# Cell Layers: Uncovering clustering structure and knowledge in unsupervised single-cell transcriptomic analysis

**DOI:** 10.1101/2020.11.29.400614

**Authors:** Andrew P. Blair, Robert K. Hu, Elie N. Farah, Neil C. Chi, Katherine S. Pollard, Pawel F. Przytycki, Irfan S. Kathiriya, Benoit G. Bruneau

## Abstract

**Motivation:** Unsupervised clustering of single-cell transcriptomics is a powerful method for identifying cell populations. Static visualization techniques for single-cell clustering only display results for a single resolution parameter. Analysts will often evaluate more than one resolution parameter, but then only report one.

**Results:** We developed Cell Layers, an interactive Sankey tool for the quantitative investigation of gene expression, coexpression, biological processes, and cluster integrity across clustering resolutions. Cell Layers enhances the interpretability of single-cell clustering by linking molecular data and cluster evaluation metrics, to provide novel insight into cell populations.

**Availability and implementation:** Upon request

## 1. Introduction

Single-cell RNA sequencing (scRNA-seq) technology allows for the classification of heterogeneous cell populations. Interpreting scRNA-seq profiles computationally requires clustering, which aims to group cells with similar transcriptional signatures (Hwang, B. *et al.* 2018). The number of clusters and subsequent cell type characterization is commonly reliant on a clustering algorithm’s parameter choice, e.g., k-means clustering. A popular scRNA-seq clustering algorithm called Louvain is parameterized by a modularity optimization technique called ‘resolution’. The resolution parameter regulates the number of clusters, with low values producing a few large clusters and higher values producing many small clusters (Traag, V.A. et al. 2019). A primary hurdle lies in providing a quantitative representation and explanation of cell type relationships in multi-resolution clustering.

Traditional scRNA-seq cluster analysis is visualized using t-distributed stochastic neighbor embedding (*t-*SNE) or uniform manifold approximation projection (UMAP) (Schwartz, G.W. *et al.* 2020). While dimensionality reduction algorithms typically provide biologically meaningful topology, they only allow users to visualize and assess one clustering resolution at a time. Furthermore, since the cluster resolution governs cell delineation, it may not subdivide all populations at a single parameter. Analysts often evaluate multiple resolutions for different analytical purposes. For these reasons, we developed Cell Layers-an interactive data visualization tool for scRNA-seq multi-resolution cluster analysis.

Cell Layers is a Sankey network. The network’s nodes are clusters. We call the edges between nodes the “flow”, which represent the transfer of cluster assignments across a cluster parameter grid search. A ternary chart can also be used for interpreting the coexpression of two to three marker genes for each flow. The Sankey representation of multi parameter clustering enables the rapid evaluation of single cell type characterization.

## 2. Methods

### 2.1. Louvain Strategies

The Louvain algorithm takes a cell-to-cell KNN graph as input. In scRNA-seq analysis, the default construction method is based on the euclidean distance of a user-defined PCA subspace (Blondel,V.D. *et al.* 2008). Cells are then iteratively grouped together to optimize the Louvain modularity function, which is thresholded by the resolution parameter. Louvain clusters are numerically labeled, with lower numbers signifying larger clusters. The output of a multi-resolution Louvain analysis is a cell by resolution parameter matrix, where values are the cluster assignment. The primary input to Cell Layers is a multi-resolution and cell by gene expression matrix.

### 2.2. Data Representation

In Cell Layers, each column of nodes represents a community structure C_vj_, where *v* represents clusters {0,1,..n} at resolution parameter 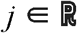 The Sankey network’s columns are ordered by a linearly increasing cluster parameter specified by a user-defined range and increment *q.* Each column’s community structure C_vj_ is represented by nodes that are scaled by the cluster sample size and arranged to minimize edge overlap. The edges or “flow” between clusters is computed as the number of cells from C_vj_ assigned to C_vj+q_. The data structure generated by Cell Layers is a directed acyclic graph.

### 2.3. Single Cell Features

The flow may be painted by marker gene expression for cell-type characterization. Users define the genes and their expression signature. To assess a single gene, each flow in the network is painted by the gene’s average expression for cells between C_vj_ to C_vj+q_ (Fig. 1A). The first drop-down menu allows users to select a gene, which will dynamically update the diagram’s flow and expression scale bar. The second drop-down menu allows the user to update node color by cluster metrics or biological process activity (BPA, see below for details). Additional drop-downs allow the user to set node width and modify the network layout. Users may choose any Matplotlib colormap for painting the flow and nodes.

**Fig. 1.**
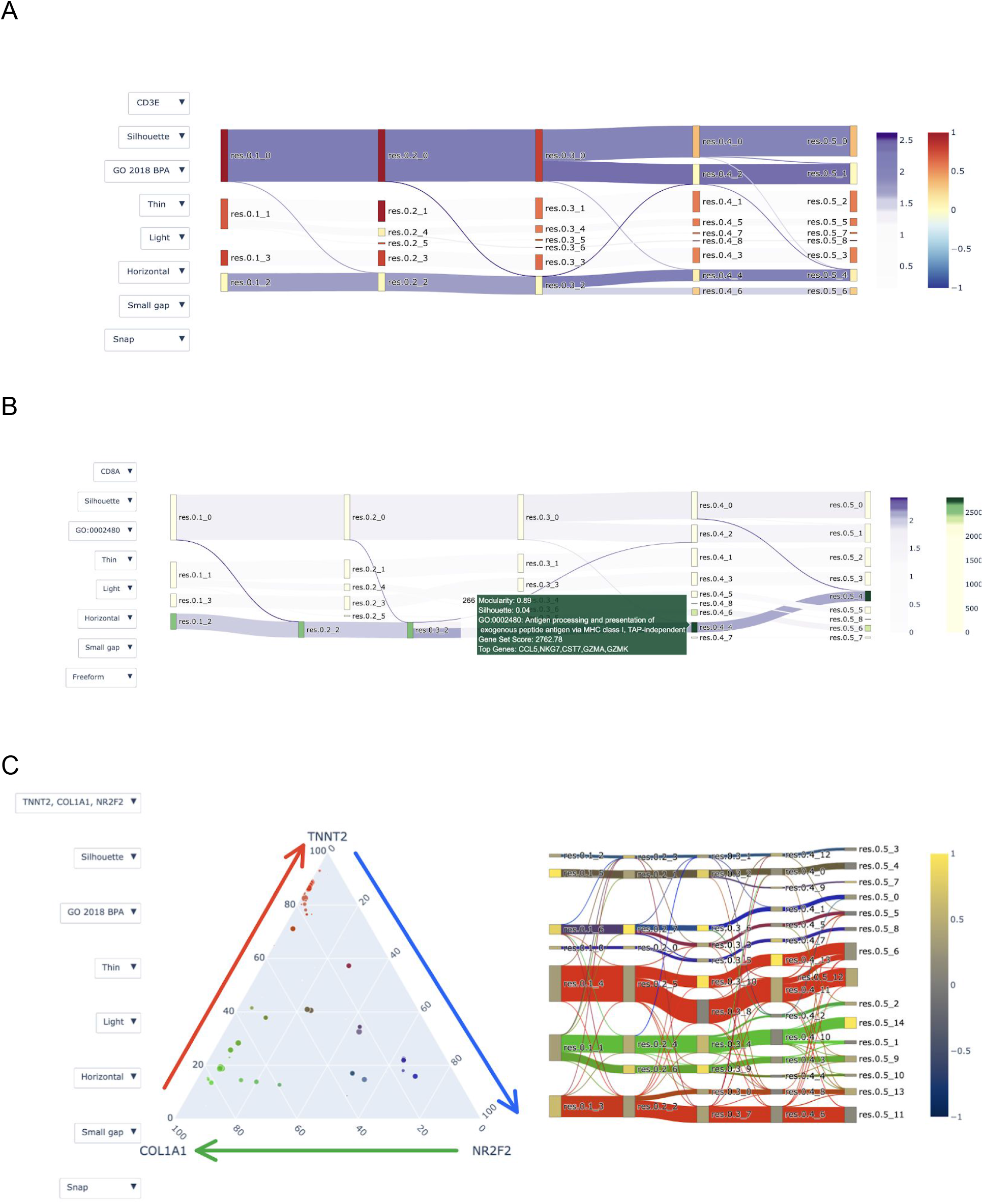
Application of Cell Layers on PBMCs or iPSC-derived cardiomyocyte differentiation (Kathiriya, I.S. et al. 2020). All nodes are labeled by their resolution parameter followed by an underscore indicating their cluster assignment. (**a**) PBMC multi resolution analysis from 0.1 to 0.5. Edges are painted by *CD3E*, which is a marker gene for CD8 T, Memory CD4 T, and Naive CD4 T cells. Nodes are painted by Silhouette score. The lower Silhouette values indicate samples are near the decision boundary of neighboring clusters. (**b**) Nodes painted by enrichR GO 2018 Biological Process gene set scores for GO:0002480. The node hover template provides users cluster performance metrics (modularity and silhouette scores), GO term title, enrichR gene set score, and the top 5 differentially expressed genes. Edges are colored by NK marker gene *CD8A.* (**c**) iPSC-derived cardiomyocyte multi resolution analysis from 0.1 to 0.5. Edges are painted by coexpression of *TNNT2* (red), *COL1A1* (green), and *NR2F2* (blue). Nodes are painted by Silhouette score. Arrows on the Ternary plot indicate the direction of the co-expression scale for each edge in the Sankey chart.

Nodes may be painted by cluster metrics or BPA. For example, to assess cluster performance users may paint a C_vj_ by their silhouette score or any cluster evaluation metric for C_vj_. Moreover, users can paint C_vj_ by BPA, which provides a robust representation of cellular states and may be used for alignment across species (Fig. 1B) (Ding,H. et al. 2019). The ability to quickly switch between cluster evaluation metrics and BPA provides users two orthogonal quantitative approaches for cell type characterization.

Many cell types are characterized by the coexpression of marker genes. To evaluate the coexpression for a user-defined geneset *E* containing *n={2,3}* genes over a flow with *m* cells, we created a gene expression percentile *(GEP)* for each gene *E*_i_. We calculated *GEP* as:

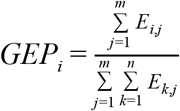

The *GEPs* for a geneset is then mapped to an RGB hex code using Matplotlib. A Plotly ternary chart is used to depict the percentile ratios of coexpressed genes for each flow (Fig. 1C). Each point in the ternary plot corresponds to a flow’s sample size and expression percentile.

### 2.4. Future Directions

Cell Layers was built for versatility and it’s application extends beyond the single cell features outlined previously. Node hover templates may include cluster metadata, which users may use to assess batch composition or integration. Downstream of multi-resolution analysis, Cell Layers can be used to evaluate data imputation models by assessing cluster stability metrics. Multi-resolution marker gene detection methods may be devised using statistical methods, such as the Jaccard Index. Additionally, protein activity profile methods could be integrated to resolve tissue-specific clusters (Ding,H. et al. 2018).

### 2.5. Software Availability

Cell Layers is integrated in the JupyterLab computing environment, which also supports popular scRNA-seq tools. Plotly is open-source software for data analytics and visualization of data science models. We made significant adaptations to Plotly’s interactive Sankey and Ternary API for scRNA-seq multi-resolution Louvain analysis. Cell Layers will be available via pip, GitHub, Docker, and Singularity.

## 3. Conclusion

Clustering of scRNA-seq data is used for computationally identifying cell populations. Analysts typically assess multiple cluster parameters, cluster performance metrics, and marker genes before annotating clusters. Fixed dimensionality reduction methods for visualizing single-cell clustering are limited by the number of attributes that users can assess. Cell Layers enables analysts to easily utilize multi-resolution parameters for interactively exploring and characterizing single cell populations.

## Supporting information

Cell Layers PBMC Example

## Acknowledgments

We thank the Cytoscape and scNetViz developers Alex Pico and Scooter Morris for their input on Plotly, Dan Carlin for his input on multi-resolution analysis, members of the CIRM Heart of Cells group, Gladstone Bioinformatics core, and Bruneau lab for discussions and comments.

## Authors Contributions

A.P.B. conceived and initiated the project. R.H and E.N.F. assisted in analysis. B.G.B., P.F.P, and I.S.K. supervised A.P.B. I.S.K. provided datasets. K.S.P. advised. All authors commented on the manuscript.

